# G protein Inactivation as a Mechanism for Addiction Treatment

**DOI:** 10.1101/2024.12.16.628727

**Authors:** Carlie Neiswanger, Micaela V. Ruiz, Kandace Kimball, Justin D. Lee, Benjamin Land, Andre Berndt, Charles Chavkin

**Affiliations:** Departments of Pharmacology, University of Washington; Seattle, WA; Bioengineering, University of Washington; Seattle, WA

**Keywords:** Addiction, kappa opioid receptor, microdosing

## Abstract

The endogenous dynorphin/kappa opioid receptor (KOR) system in the brain mediates the dysphoric effects of stress, and KOR antagonists may have therapeutic potential for the treatment of drug addiction, depression, and psychosis. One class of KOR antagonists, the long-acting norBNI-like antagonists, have been suggested to act by causing KOR inactivation through a cJun-kinase mechanism rather than by competitive inhibition. In this study, we screened for other opioid ligands that might produce norBNI-like KOR inactivation and found that nalfurafine (a G-biased KOR agonist) and nalmefene (a KOR partial agonist) also produce long-lasting KOR inactivation. Neither nalfurafine nor nalmefene are completely selective KOR ligands, but KOR inactivation was observed at doses 10-100 fold lower than necessary for mu opioid receptor actions. Daily microdosing with nalfurafine or nalmefene blocked KORs responsible for antinociceptive effects, blocked KORs mediating stress-induced aversion, and mitigated the aversion during acute and protracted withdrawal in fentanyl-dependent mice. Both nalfurafine and nalmefene have long histories of safety and use in humans and could potentially be repurposed for the treatment of dynorphin-mediated stress disorders.

## Introduction

An essential role for endogenous dynorphin opioids in regulating mood and addiction risk has been suggested by multiple preclinical studies during the last two decades (1–4).

Pharmacological and genetic disruption of dynorphin actions in the rodent brain block depression-like and anxiety-like behaviors (5–7). Stress-induced release of endogenous dynorphins potentiates the rewarding effects of addictive drugs (e.g. cocaine, ethanol, nicotine, and heroin) (8–12). Stress-induced reinstatement of drug self-administration is blocked by kappa opioid receptor (KOR) antagonists (13), and KOR antagonists block the dysphoric effects evident during opioid, nicotine, and ethanol abstinence (1, 14, 15). Kappa agonists have psychotomimetic effects (16), and the stress-induced release of dynorphins disrupts prefrontal-cortical circuits responsible for cognition (17, 18). Stress exposure has been shown to increase the risk of mood and substance use disorders in humans (19, 20), and these preclinical studies strongly suggest that KOR antagonists could be therapeutically useful in promoting stress resilience in vulnerable people. However selective, safe, and effective KOR antagonists for human studies are still being developed, and the role of dynorphin in human pathophysiology is not yet established.

The majority of the preclinical studies examining the dyn/KOR system have used the selective KOR antagonists norbinaltorphimine (norBNI) or JDTic (15, 21). Although these drugs are competitive KOR antagonists in vitro, they both have the unusual properties of delayed onset of action and very long durations of effect when given in vivo. For example, a single injection of norBNI in rodents or nonhuman primates can block KOR activation for weeks (22, 23). The mechanism for this long duration was initially unclear and attributed to a very slow clearance of the drug from the brain (a pharmacokinetic mechanism) (24). However, a detailed analysis of the molecular pharmacology revealed that both norBNI and JDTic are not conventional antagonists; rather they are functionally selective KOR agonists that activate cJun N-terminal Kinase (JNK) (22, 25), which recruits the phospholipase peroxiredoxin 6 (PRDX6) to the plasma membrane; PRDX6 bound to KOR stimulates NADPH oxidase to locally generate reactive oxygen (ROS) which oxidizes the sulfhydryl in exposed cysteine residues and depalmitoylates the G-protein, Gɑi. The depalmitoylated Gɑi binds more tightly to KOR and prevents receptor activation (26, 27). Pharmacological inhibition of the JNK or PRDX6 components in this signaling pathway or genetic deletion of *JNK-1* blocked the long duration of norBNI action (22, 25, 27). In this context, the agonist and antagonist labels are misleading, and norBNI-like ligands are better described as *KOR-inactivating agents*.

This series of molecular characterization studies helped establish that there are four different types of functionally selective KOR ligands (Fig. 1). These include unbiased high-efficacy agonists, competitive antagonists, and long-acting KOR-inactivating agents. A fourth group of functionally selective KOR ligands includes nalfurafine, a highly efficacious but G-biased KOR agonist (28, 29), and nalmefene, a low-efficacy KOR partial agonist (30, 31) (Fig. 1D).

**Figure 1:**
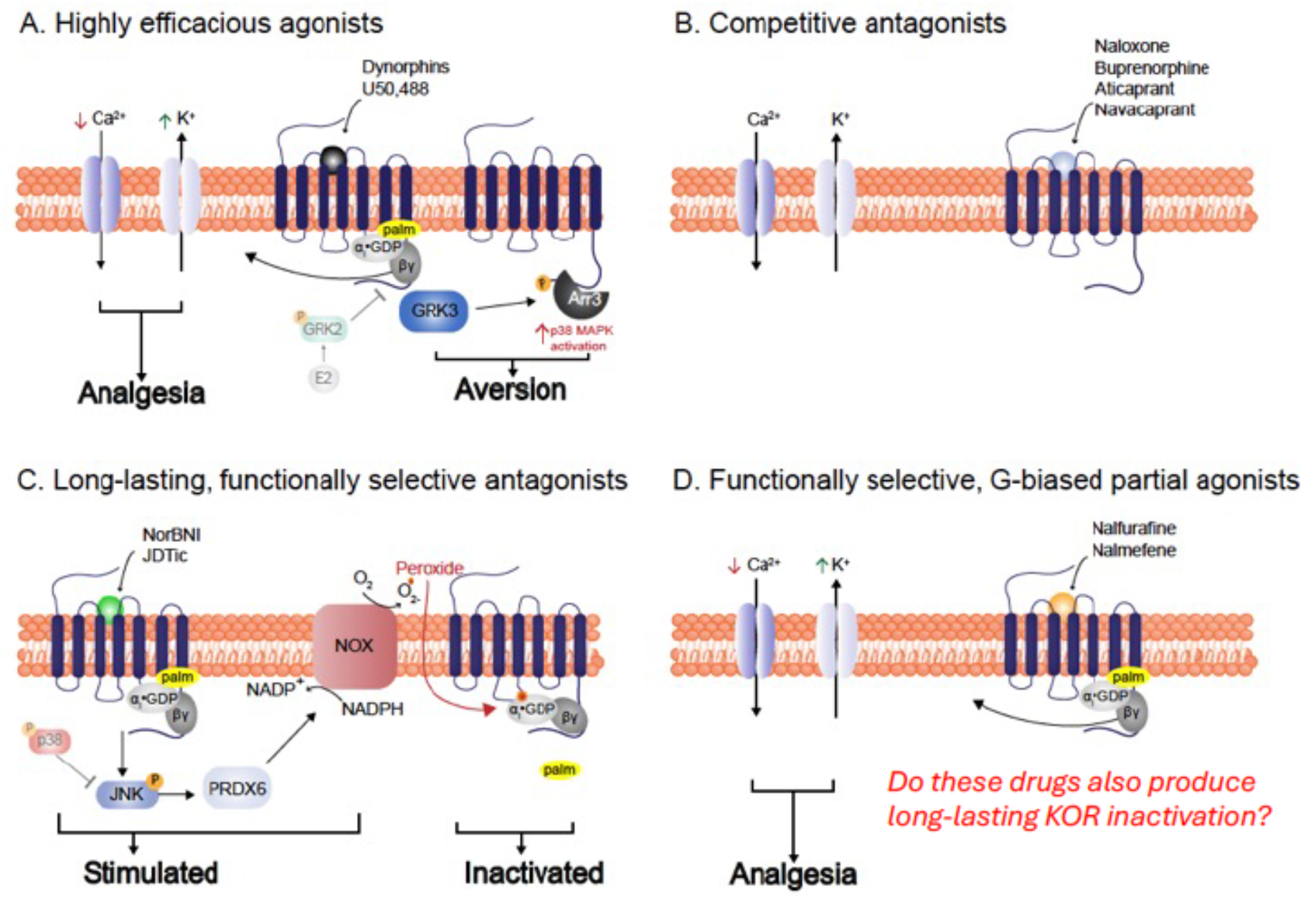
An overview of major pathways in the functionally selective signaling cascade of the kappa opioid receptor. (**A**) Highly efficacious agonists including U50,488 and the endogenous dynorphin peptides activate both heterotrimeric G-proteins and β-arrestin- dependent signaling cascades. G-protein activation releases Gβγ that activates inwardly rectifying potassium channels and inhibits voltage-gated calcium channels to reduce neuronal excitability and presynaptically inhibit neurotransmitter release. The dysphoric effects of efficacious KOR agonists occur through GRK3/β-arrestin activation of p38α MAPK in serotonergic neurons in the dorsal raphe nucleus (57, 58) and dopaminergic neurons in the ventral tegmental area (59). (**B**) Competitive antagonists including naloxone bind to the orthosteric site of the receptor and sterically inhibit agonist binding. (**C**) Long–lasting, functional antagonists including norBNI are proposed to be functionally selective activators of a signaling cascade that leads to the phosphorylation of c-Jun N-terminal kinase (JNK), recruitment of PRDX6, increased activity of NADPH oxidase, and generation of reactive oxygen species (ROS) (27). Oxidation is proposed to depalmitoylate Gαi and shift the orientation of the receptor•G-protein binding interface to irreversibly block guanine nucleotide exchange. Because activated p38α MAPK inhibits JNK, highly efficacious KOR do not cause long-lasting KOR inactivation (26). (**D**) Biased and partial agonists cause conformational changes that lead to the G-protein pathway being preferentially activated (nalfurafine) or the GRK3/β-arrestin pathway being activated weakly (nalmefene) and not causing aversion. The present study tests the prediction that KOR partial agonists that do not efficiently activate GRK3/β-arrestin signaling will inactivate KOR through a norBNI-like mechanism.

Because neither nalfurafine nor nalmefene efficiently activates GRK3/β−arrestin-dependent desensitization mechanisms (32, 33), we hypothesized that they might activate JNK/PRDX6 to produce a norBNI-like long-lasting inactivation of KOR. Nalfurafine and nalmefene have long histories of safe use in humans (34, 35) and could conceivably be repurposed as medications to block KOR and promote stress resilience for the adjunctive treatment of mood disorders, psychosis, and drug addiction.

## Materials and Methods

### Animals

Adult C57BL/6 wild-type (WT) or KOR-cre male and female mice were used (Jackson Labs). Animal studies were approved by the University of Washington IACUC.

### Drugs

Nalfurafine (NIDA Drug Supply Program), nalmefene (Tocris), norBNI (NIDA Drug Supply Program), U50,488 (Tocris), naloxone (Tocris), naltrexone (Tocris), GNTI (Tocris), and MJ33 (Sigma Aldrich) were dissolved in saline and administered intraperitoneally (i.p.). Estradiol (50 ug/kg; Cayman Chemical) and progesterone (5 mg/kg; ThermoFisher) were dissolved in 0.1% ethanol/ 0.1% Cremophor EL/ 99% saline and administered subcutaneously (s.c.) in a volume of 10ml/kg.

### Stereotaxic injection

Male KOR-cre mice between 5-7 weeks were anesthetized with 2.5% isoflurane and head fixed on a Kopf Model 942 stereotaxic alignment system. A 2.0 mL model 7002 KH Neuros Syringe (Hamilton, NV, USA) was lowered for viral injection of 0.5 mL of AAV1-FLEX-oROS- mCherry. For in vitro brain slice experiments, bilateral injection into the VTA using coordinates: ML: +/- 0.5 mm, AP: −3.28 mm, DN: −4.5mm, and for fiber photometry experiments, unilateral injection using coordinates AP: −3.28 mm, ML: -1.71 mm, DV: −4.67 mm at a 15-degree angle; injections were done 2-4 weeks prior to imaging. An implantable fiber optic cannula (400/430 core, 0.57 NA; Doric Lenses, Quebec, CA) was placed at the same coordinates as the viral injection site, and then C&B Metabond was used to secure the cannula to the skull. After injection, the needle was kept at the injection site for 5 minutes before removal. All mice were monitored and given 10 mg/kg carprofen daily for 7 days for post-surgical analgesia.

### Fiber photometry

Briefly, a real-time signal processor (RZ5P; Tucker-Davis Technologies) was connected to Synapse Software (Fiber Photometry) to set the frequency of light stimulation and record input from photodetectors as described (43). For in vivo experiments, mice were run with randomized treatments on separate days. Mice received saline, nalfurafine (100 µg/kg), or nalfurafine (100 µg/kg) with naloxone (10 mg/kg). For 100 µg/kg nalfurafine experiments, after baseline, the recording session lasted for 45 minutes. For 10 mg/kg naloxone pretreatment to 100 µg/kg nalfurafine experiments, after baseline, mice were given administration of naloxone, and after 15 minutes 100 µg/kg nalfurafine was administered. Mice were allowed to freely explore the chamber during in vivo recordings. N = 3-13 per treatment group.

### Estrous cycle stage determination

Vaginal lavage for the estrous cycle stage determination was done following behavioral testing; 50 mL of deionized water was pipetted to collect epithelial cells and placed on glass slides for cytology. The estrous cycle stage was classified using the estrous cycle identification tool (56) and then quantified by the relative ratio of nucleated epithelial cells, cornified epithelial cells, and leukocytes. Cytology was determined with the investigator blind to treatment.

### Hormone treatment

Female mice were injected subcutaneously with 5mg/kg progesterone (ThermoFisher) or 50ug/kg estradiol (Cayman Chemical). 3 days later, the mice were tethered for fiber photometry (as described above) and given an intraperitoneal injection of nalfurafine (100 µg/kg), and the oROS response was measured. Estrous states were determined after the recording session. Following lavage, mice were given a second subcutaneous injection of the treatment they received prior and 3 days later, repeated the assessment of their estrous state and fiber photometry experiments. Both treatment groups had a two-week recovery period with no treatment. Following the recovery period, the treatments were switched. Data analyses were done by an investigator blind to treatment. N = 7-12 per treatment group.

### In vitro neuron imaging

Brains were collected 2-4 weeks after viral injection, and 200 µm horizontal slices were obtained using a vibratome as previously described (39). Image collection was done on a Bruker Investigator 2-photon microscope, with software Prairie View 5.5, simultaneously collecting the mCherry (1040 nm fixed) and GFP (920 nm tunable) signals with a Nikon 16X water immersion objective, as well as a z-stack spanning 60 µm across an hour time course. During this collection time, a baseline was established for 7 minutes in ACSF buffer solution (124 mM NaCl, 3 mM KCl, 2 mM MgSO_4_, 1.25 mM NaH_2_PO_4_, 2.5 mM CaCl_2_, 26 mM NaHCO_3_, 10 mM Glucose) or ACSF with naloxone (10 µM), MJ33 (10 µM), or JNK-IN-8 (1 µM) at 32°C followed by treatment with nalfurafine (100 nM), nalmefene (10 µM), or those in addition to previously mentioned inhibitors for 30 min, followed by H_2_O_2_ (590 µM) for 5 minutes. Analyses were completed by creating a hyperstack in ImageJ and then creating a Z-projection for both signal channels. Regions of interest (ROIs) were drawn broadly over the entire slice, and values were collected using ImageJ for each channel with the same ROI from raw, unaltered images. The mCherry signal value was subtracted from the GFP signal value, and the resulting data were fit so that the lowest value was adjusted to 1. Then ΔF/F was calculated using the average value of the baseline (7 minutes of buffer solution, or 7 minutes of only inhibitor treatment) for the rest of the time course. Cells that showed no response to H_2_O_2_ wash were excluded from the analysis. Representative 2p images (Fig 2.) have enhanced contrast for visual representation. N = 3-10 per treatment group, with each data point representing the average of 15-30 individual cells in a single slice.

**Figure 2:**
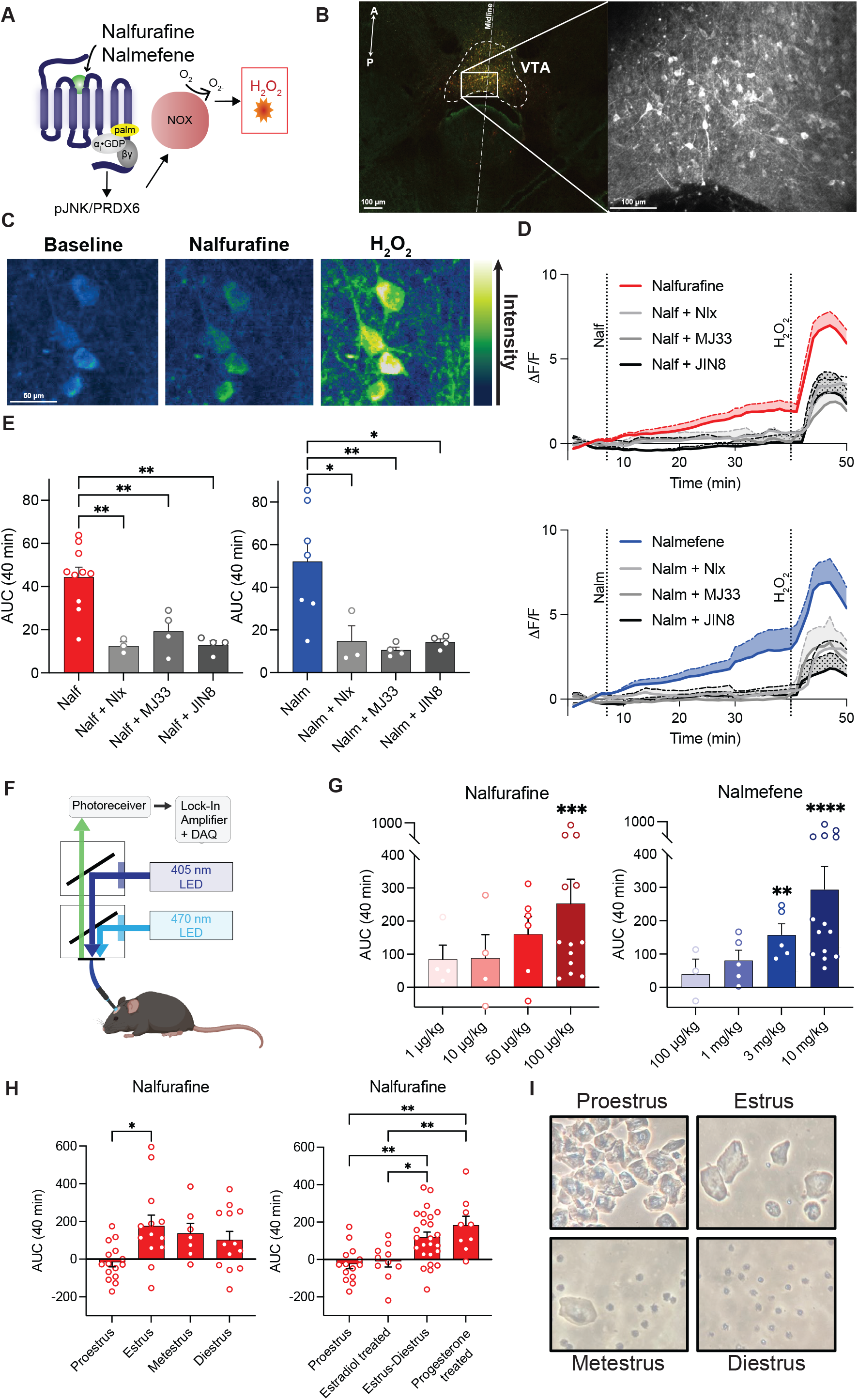
Nalfurafine and nalmefene cause ROS generation in a KOR-specific manner. (**A**) Proposed pathway for nalfurafine and nalmefene leading to the generation of ROS. (**B**) *Left* Confocal image of a horizontal VTA slice showing merged channels of the GFP sensor (green) and the mCherry tag (red). *Right* Representative 2-photon image of the same live slice at higher magnification. (**C**) Representative images (contrast-enhanced) using 2-photon microscopy of KOR-positive cell bodies in the VTA with oROS expression under buffer-only condition (baseline), after 100 nM nalfurafine washed on for 20 min, followed by H_2_O_2_ for positive control. (**D**) ΔF/F values for 50 min imaging sessions. Wash on of 100 nM nalfurafine (red, *N* = 10) and 10 μM nalmefene (blue, *N = 7*) both show clear increases in oROS fluorescence that can be blocked by naloxone (*N* = 3), MJ33 (*N* = 4), or JNK-IN-8 (*N* = 4). The addition of H_2_O_2_ at 40 min confirmed that the sensor was active at the end of the time course. (**E**) AUC quantification of time course data from 0-40 min. (**F**) Visual of the fiber photometry setup used in (G). (**G**) AUC quantification of fiber photometry time course for 40 min after injection of various doses of nalfurafine (*N* = 4-13) and nalmefene (*N* = 3-14) which dose- dependently increased oROS fluorescence. (**H**) (Left) AUC quantification of time course data after injection of 100 µg/kg nalfurafine in female mice measured at various points in the estrus cycle (*N* = 7-16). Female mice show no response to nalfurafine at the high estrogen stage (proestrus). (Right) AUC 0-40 min of female mice pretreated with estradiol or progesterone compared to the cycle collapsed into high estrogen stage (proestrus) and low estrogen stages (estrus-diestrus) (*N* = 9-16). Pretreatment with estradiol mimics the proestrus stage, and pretreatment with progesterone keeps the animals in the low estrus state, mimicking the low estrogen stages. (**I**) Representative images of histology from vaginal lavage to determine the estrous cycle stage. Statistical comparisons are shown * p <0.05, ** p <0.01, *** p <0.001, **** p < 0.0001

### Confocal images

Horizontal images were collected using slices that underwent 2Photon imaging and were put into 10% formalin. Confocal images are taken with a Leica SP8x Confocal microscope.

### Warm water tail withdrawal

Mice were tested for U50,488 (10 mg/kg i.p.) antinociception using a 52.5°C warm water tail withdrawal test as previously described (22). Scores were calculated as the difference between the before and 30 min after U50,488 injection. N = 6-16 per treatment group.

### Odorant-paired repeated forced swim stress

Mice were treated with saline, norBNI, nalfurafine, or nalmefene daily for 7 days. To allow drug clearance and focus on the long-lasting KOR inactivation, 7 days after their last treatment mice were exposed to one 15-minute swim on day 1 and four 6-minute swims on day 2, in 30°C water, without opportunity for escape. This repeated swim stress protocol was previously demonstrated to evoke dynorphin release (8). Odorant-swim stress pairing was conditioned as previously described (42). N = 6-16 per treatment group

### Odorant-paired precipitated fentanyl withdrawal

Mice were treated with saline, norBNI, nalfurafine, or nalmefene daily for 7 days. On day 4 of the 7-day treatment, osmotic pumps (Alzet Model 1007D) filled with either saline or fentanyl (2 mg/kg/day) were implanted under the skin between the scapula to induce opioid dependence. The pumps were removed after 7 days and mice were allowed to recover for 2 days. On either day 3 (acute withdrawal) or day 10 (protracted withdrawal) post-pump removal, they were given injections of 1 mg/kg naloxone in the presence of a Nestlet containing 20 µl of imitation almond extract once in the morning and once in the afternoon. Odorant-conditioned aversion was assessed in the 3-compartment place preference apparatus 7 days after naloxone- odorant pairing.

### Odorant-aversion test

Mice underwent two types of odorant-conditioned pairing: stress-odorant pairing in which almond scent was paired with repeated forced swim stress and withdrawal-odorant pairing in which almond scent was paired with two instances of naloxone-precipitated withdrawal (outlined above). Mice were pre-exposed to a 3-chamber Plexiglas box for 3 min before odorant conditioning. For the odorant-aversion test, mice were placed in a Plexiglas 3-chamber box with a quarter Nestlet containing 20 µl of almond extract placed on one far side of the chamber, and a quarter Nestlet with no scent placed on the far side of the other chamber. The session in the 3-chamber box was video recorded for 14 min and analyzed using Ethovision for the time spent in each zone. The odorant aversion was calculated by the time spent in the odor-paired chamber minus the time spent in the opposite chamber. N = 8-15 per treatment group.

### Prolactin determination

Blood samples of up to 500 µL were taken from mice 1 hour after drug treatment and allowed to clot for 5 min at room temperature before being centrifuged for 15 min (2000 x g) at 4°C. Serum was immediately frozen on crushed dry ice and stored at -80°C until assayed in Mouse Prolactin (PRL) ELISA Kit PicoKine® from Boster Biological Technology. PRL assay was performed following the manufacturer’s instructions at 1:10 dilution of sera. Contrary to the instructions, we found that sera reduced the slope of the PRL standard curve. We added 10 µL of pooled sera from control, uninjected mice to each 100 µL of diluted PRL standard to correct for this. Thus, sera PRL assay values reported are ng/ml over baseline. N = 4-16 per treatment group.

### Diuresis assay

Mice were habituated for 20 min in 4x4 inch plexiglass boxes with pre-weighed paper towels below them before being injected with saline or 100 ug/kg nalfurafine. 60 min after injection, mice were removed, and the change in weight of the paper towel was measured. Mice were also tested 1 and 24 hours, 7- and 14-days post 10 mg/kg norBNI treatment or 7 days after the last administration of 7 daily treatments of 1 mg/kg norBNI. For the 30-day study, mice were injected with either saline or nalfurafine before urine output measurement every day for 30 days. N = 10-20 per treatment group and N = 3-4 for 30-day study.

### GNTI-induced pruritus

Mice were injected with saline (s.c.) 20 min prior to 30 µg/kg GNTI (s.c.), and scratching behavior was recorded for 30 min as described (29). Mice were then given daily treatments of either saline or nalfurafine (50 µg/kg) for 7 days. The following day, mice were injected with nalfurafine (50 µg/kg) 20 min prior to 30 µg/kg GNTI to determine nalfurafine block of scratching behavior.

### Statistical analysis

All data were analyzed using either one- or two-way ANOVA appropriate for variables using GraphPad Prism unless otherwise stated. Post-hoc tests for multiple comparisons were performed to determine differences from controls. Initial analysis was done with experimenters blinded to treatment groups. Exclusion criteria for all experiments were based on statistical outliers or insufficient responses during baseline measures or compared to positive or negative controls. The number of animals used in each treatment group was determined by the prior data for similar assays. Animals in each study were age-matched and randomized before the start of treatments. Studies were designed so that all animals in a single cage received the same treatment as their cage mates.

## Results

### The novel sensor oROS-Gr detects increases in ROS generation in the presence of nalfurafine and nalmefene

Because the JNK/PRDX6 mechanism for receptor inactivation is dependent on the generation of ROS, we reasoned that monitoring ROS generation would be an effective means to identify compounds having norBNI-like effects. To achieve this, we optimized a ROS sensor using a structure-based design (36). The resulting sensor oROS-Gr performed better than previous iterations (Fig. S1) and was expressed in a cre-dependent manner in the ventral tegmental area (VTA) of KOR^Cre^ mice (17). Ex vivo horizontal slices were used for live-cell 2-photon fluorescence imaging (Fig. 2B). Consistent with our hypothesis that the G-biased KOR agonists nalfurafine and nalmefene would produce long-lasting norBNI-like effects, we observed that nalfurafine (100 nM) and nalmefene (10 μM) significantly increased oROS fluorescence (Fig. 2D-E). The fluorescence increase was blocked by the competitive opioid receptor antagonist naloxone (10 μM), the PRDX6 inhibitor MJ33 (10 μM) (37), and the JNK inhibitor JNK-IN-8 (1 μM) (Fig 2D). Activation of oROS-Gr in the VTA was also imaged using in vivo fiber photometry (Fig 2F). Both nalfurafine and nalmefene caused dose-dependent increases in fluorescence, and the response was blocked by pretreatment with 10 mg/kg naloxone (Fig. 2G). These results confirm that both nalfurafine and nalmefene stimulate the generation of ROS through the KOR/JNK/PRDX6 signaling cascade in KOR-expressing VTA neurons.

### KOR-induced ROS generation increase is estrus cycle dependent in female mice

The generation of oROS-Gr allowed us to address an additional fundamental question relating to the regulation of KOR sensitivity to the long-lasting effects of norBNI. It has been previously reported that female rodents have estrus state-dependent insensitivity to KOR activation because estrogen stimulates G protein Receptor Kinase 2 (GRK2) to block Gβγ signaling (38). Through this proposed mechanism, estrogen blocks the long-lasting effects of norBNI (38, 39). To determine if the activation of oROS was similarly affected, we expressed oROS-Gr in VTA neurons of adult female KOR-Cre mice and measured the fluorescence responses to nalfurafine during different estrus stages. Nalfurafine significantly increased oROS fluorescence during estrus (lowest estrogen state) but had no effect during proestrus (highest estrogen state) (Fig. 2H). Female mice pretreated with estradiol had no significant oROS response to nalfurafine, whereas female mice pretreated with progesterone showed a robust increase in fluorescence after nalfurafine injection (Fig. 2I). These results are consistent with previously suggested mechanisms of inconsistent G protein-mediated KOR agonist effects in females.

### Repeated administration of nalfurafine or nalmefene block kappa agonist-induced analgesia in a JNK-pathway-specific manner

A single injection of 10 mg/kg norBNI completely inactivates KOR for weeks in mice (22). A single injection of a 100-fold lower dose of norBNI was ineffective, but daily injection of 0.1 mg/kg norBNI for 3 weeks produced KOR inactivation in both male and female mice (40). Presumably, the lower dose inactivates a small percentage of the receptors each day, and because KOR recovery takes weeks, receptor inactivation accumulates. Reducing a drug dose by 100-fold reduces the risk of adverse and off-target effects and would be potentially advantageous in human therapeutics. To determine the abilities of nalfurafine and nalmefene to produce norBNI-like cumulative KOR inactivation, we injected various doses daily for 7 days and then waited for 7 days to allow drug clearance before assessing the degree of KOR inactivation using the standard warm-water (52.5°C) tail withdrawal assay. Treatment for 7 days with low doses of norBNI (330 µg/kg), nalfurafine (100 ng/kg), or nalmefene (100 µg/kg) significantly inhibited the antinociceptive effects of the KOR agonist U50,488 (Fig 3A-C). The inactivation was long-lasting: norBNI-treated mice (1 mg/kg) showed only partial recovery after 28 days (Fig 3D). Nalfurafine-treated mice recovered only after 21 days (Fig 3E), and nalmefene-treated mice required more than 14 days to recover (Fig 3F). In contrast, the effects of naloxone (10 mg/kg) and naltrexone (10 mg/kg) completely reversed in 3 days (Fig 3 G-H). When PRDX6 was inhibited by injection with MJ33 (1.25 mg/kg) before each injection of norBNI, nalfurafine, or nalmefene, the rates of recovery of the U50488 analgesic response were significantly increased. The sensitivity to the selective PRDX6 inhibitor supports the hypothesis that long-acting KOR inactivation measured in vivo results from the JNK/PRDX6/ROS mechanism proposed. The difference between naltrexone and nalmefene in their durations of action is striking since these two drugs have very similar structures (a ketone versus a methylene group on ring carbon 6). These results support the prediction that microdosing with nalfurafine or nalmefene can inactivate KOR in a norBNI-like manner through the JNK/PRDX6 cascade.

**Figure 3:**
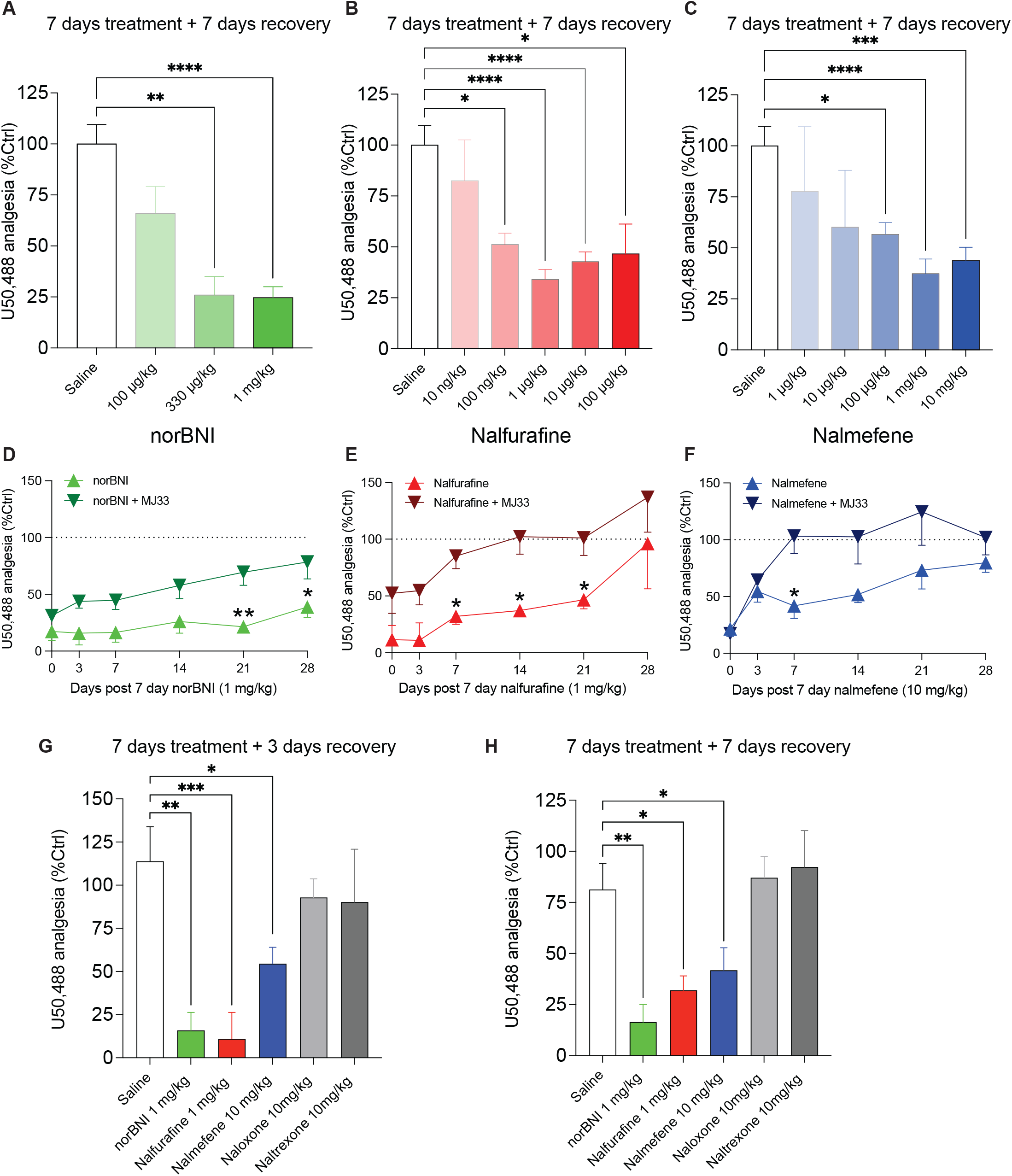
Repeated administration of nalfurafine and nalmefene blocks KOR-induced analgesic response through the JNK pathway. Tail withdrawal latency is shown as a percent of the baseline of the normal increase elicited by U50,488 administration. (**A-C)** 7 days post-last treatment norBNI (100 µg/kg - 1 mg/kg, *N* = 4- 20), nalfurafine (10 ng/kg - 100 µg/kg, *N* = 7-24), and nalmefene (1 µg/kg - 10 mg/kg, *N* = 8- 24). All 3 treatments dose-dependently block U50,488-induced analgesia. (**D-F)** The same dosing regimen as in A-C. Treatments of norBNI (1 mg/kg, *N* = 8), nalfurafine (1 mg/kg, *N* = 11), and nalmefene (10 mg/kg, *N* = 14) were given once daily for 7 days. Time points are shown as days after the last administration of treatment measured out to 28 days. Repeated treatment with norBNI, nalfurafine, and nalmefene blocks U50,488-induced latency increase. Pretreating with MJ33 1 hour before each daily dose caused KOR function to recover more rapidly. (**G)** 3 and (**H)** 7 days post-last treatment of 7 daily doses of Saline (*N* = 16), norBNI (1 mg/kg, *N* = 8), nalfurafine (1 mg/kg, *N* = 11), nalmefene (10 mg/kg, *N* = 14), naloxone (10 mg/kg, *N* = 7), and naltrexone (10 mg/kg, *N* = 8). Statistical comparisons are shown * p <0.05, ** p <0.01, *** p <0.001, **** p < 0.0001

### Repeated administration of nalfurafine and nalmefene block stress-paired cue aversion

To determine if microdosing nalfurafine or nalmefene would block dynorphin-mediated stress responses, we used two different assays of dysphoria. In mice, dysphoria is typically measured as aversion conditioned to a cue associated with the stressful experience (41). In the first assay, mice were subjected to a repeated forced swim paradigm previously shown to release dynorphin (8, 42) in the presence of a neutral odorant cue (almond scent). Following conditioning, mice were introduced to a 3-chamber place preference apparatus with the almond scent present in only one compartment (Fig. 4D). Control mice exposed to the almond scent in the absence of the stress-pairing showed no aversion to the almond scent-containing compartment. In contrast, mice experiencing stress-pairing show robust odorant-aversion (Fig. 4A-C). Pretreatment of the mice for 7 days with norBNI (either 330 µg/kg or 1 mg/kg), nalfurafine (1 µg/kg or 10 µg/kg), or nalmefene (100 µg/kg or 1 mg/kg) to inactivate KOR, blocked odorant aversion. The drug potencies in the swim-odorant assay were comparable to the tail-flick analgesia assay (Fig. 3). Because the place aversion assay requires conditioning, it is important to note that norBNI does not affect learning. In previous work, mice that were injected with cocaine in the presence of the almond scent developed robust place preference for the almond-scent paired compartment. NorBNI (10 mg/kg) treatment before cocaine conditioning did not block the development of almond-scent preference (42).

**Figure 4:**
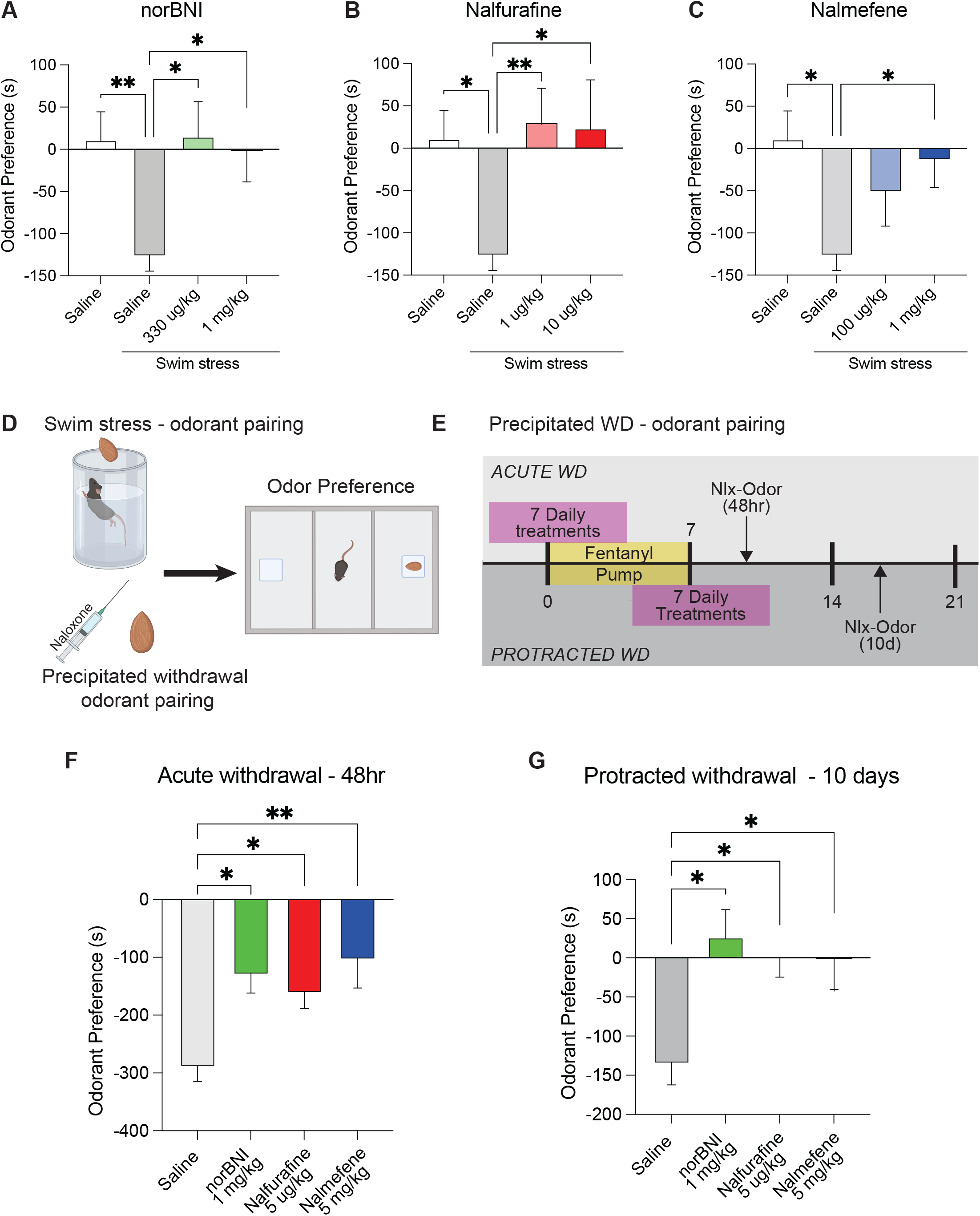
Repeated administration of nalfurafine and nalmefene block stress-paired cue aversion. (**A-C**) Odorant scores were determined by the amount of time spent in the null odorant side subtracted from the paired odorant side (a negative score representing aversion). Unstressed animals (*N* = 10) had neutral scores while stressed animals pretreated with saline (*N* = 13) showed an aversion. 7 daily injections of norBNI (330 µg/kg (*N* = 7) or 1 mg/kg (*N* = 11)), nalfurafine (1 µg/kg (*N* = 11) or 10 µg/kg (*N* = 6)), and nalmefene (1 mg/kg (*N* = 12)) blocked stress-odor paired aversion. (**D**) Schematic of almond pairing with either repeated force swim-stress or naloxone precipitated withdrawal, followed by 3-chamber aversion assay with the scent on one side. (**E**) A timeline illustrating differences between acute and protracted withdrawal groups in the naloxone-precipitated withdrawal-odorant pairing and testing. (**F**) Mice that underwent precipitated withdrawal during the acute withdrawal phase (2 days after pump removal) and were treated daily for 7 days with saline (*N* = 10) showed a strong aversion to the almond scent which was significantly attenuated in groups receiving daily injections of norBNI (*N* = 8), nalfurafine (*N* = 10), and nalmefene (*N* = 10). (**G**) Mice that underwent precipitated withdrawal during the protracted withdrawal phase (10 days after pump removal) and were treated daily for 7 days with saline (*N* = 10) showed a strong aversion to the almond scent which was fully blocked in groups receiving daily injections of norBNI (*N* = 10), nalfurafine (*N* = 10), and nalmefene (*N* = 15). Statistical comparisons are shown * p <0.05, ** p <0.01

In the second stress assay, we paired almond scent with naloxone precipitated withdrawal in fentanyl-dependent mice (Fig. 4D,E). The 1 mg/kg dose of naloxone chosen will block mu-opioid receptors but is insufficient to block KOR (43, 44). Acute opioid abstinence causes dynorphin release and includes a profound dysphoric response (43). Mice were implanted with osmotic minipumps containing fentanyl (2 mg/kg/day) to produce opioid dependence. After 7 days, the pumps were removed and 2 days later mice were injected twice with 1 mg/kg naloxone in the presence of the almond scent, once in the morning and once in the evening to precipitate withdrawal. Fentanyl-dependent mice developed a robust aversion to the odorant (Fig 4F). In contrast, control mice with saline-filled minipumps did not develop aversion when injected with naloxone in the presence of almond scent. Fentanyl-dependent mice that were pretreated for 7 days with norBNI (1 mg/kg) showed significantly reduced odorant aversion when challenged with naloxone 2-days after pump removal (acute withdrawal). Mice pretreated for 7 days with nalfurafine (5 µg/kg) or nalmefene (5 mg/kg) showed significantly reduced aversion, but the decrease was incomplete (Fig 4F). Mice that were pretreated for 7 days with norBNI (1 mg/kg), nalfurafine (5 µg/kg), or nalmefene (5 mg/kg) and experienced naloxone-precipitated withdrawal 10 days after pump removal (protracted withdrawal) showed no aversion to the odorant (Fig 4G). These results from both stress assays suggest that repeated low-dose treatment with nalfurafine or nalmefene promotes stress resilience and supports their potential therapeutic utilities in preventing relapse during protracted opioid abstinence.

### Long-term KOR antagonism is tissue-specific

The long-lasting inactivation of KOR that results from ROS generation and subsequent receptor inactivation has been previously shown to have tissue selectivity; norBNI inactivates KOR located on VTA somatic membranes but not on nerve terminals of the same neurons (39). We next determined the regulation of KOR controlling pituitary function, diuresis, and itch behaviors (Fig. 5). Nalfurafine, nalmefene, and U50,488 each dose-dependently increase serum prolactin (Fig. 5A). As expected, acute nalfurafine is a high potency, full agonist (EC50 = 2.9 µg/kg) and nalmefene is a low potency partial agonist in this assay (Fig. 5A). Pretreatment with norBNI (10 mg/kg, 7 days prior) blocked the increase in prolactin seen with U50,488 and nalfurafine (Fig. 5A). Similarly, 7 days of treatment with nalfurafine or nalmefene dose-dependently blocked the increase in serum prolactin stimulated by an acute challenge with nalfurafine (100 μg/kg) (Fig. 5B-C).

**Figure 5:**
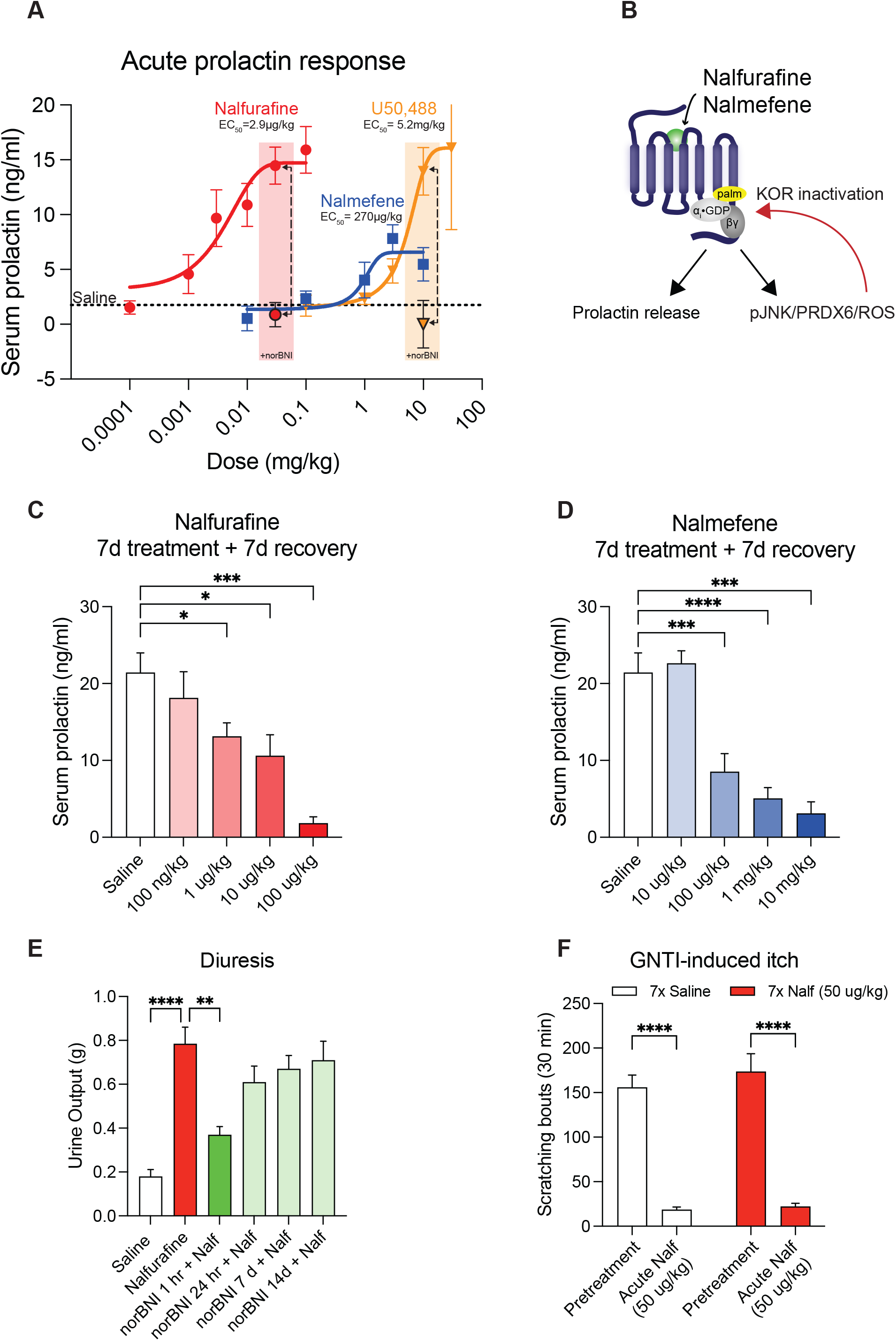
KOR inactivation demonstrates tissue specificity. (**A**) Serum prolactin dose-response curves 60 min post administration of nalfurafine (*N* = 4-12), nalmefene (*N* = 4-8), or U50,488 (*N* = 4-11). Nalfurafine has greater potency than U50,488, while nalmefene has similar potency but reduced efficacy, highlighting its partial agonism. (**B**) KOR activation by nalfurafine or nalmefene acutely increases prolactin, whereas repeated doses cause inactivation of the receptor (**C-D**) Repeated doses of nalfurafine (*N* = 4-11) and nalmefene (*N* = 4-11) blocked the nalfurafine-induced increase in serum prolactin 7 days after last treatment at various doses. (**E**) Nalfurafine (100 µg/kg) increases diuresis, which can be blocked acutely, but not long-term, by norBNI (10 mg/kg) (*N* = 10-19). (**F**) GNTI (30 ug/kg) induces pruritus, which can be blocked by nalfurafine (50 µg/kg). Daily doses of nalfurafine (7x 50 ug/kg, red bars) do not prevent this effect (*N* = 7). Statistical comparisons are shown * p <0.05, ** p <0.01, *** p <0.001, **** p < 0.0001

Kappa agonists promote diuresis, and 100 μg/kg nalfurafine increases urine output (Fig. 5E). Acute norBNI (10 mg/kg) blocked nalfurafine-induced diuresis at 1hr and 24hr post-administration but norBNI antagonism was not evident 7- or 14-days post-administration (Fig. 5E). Nalfurafine stimulation of urine output was dose-dependent, but norBNI (1 mg/kg) treatment for 7 days did not shift the dose-response curve (Fig. S2). Daily injection of 100 μg/kg nalfurafine for 30 days produced a consistent increase in urine output with no evidence of tolerance (i.e., KOR inactivation) (Fig. S2).

It has also been shown that nalfurafine can block GNTI-induced pruritis in a KOR-dependent manner (29, 45). After 7 days of daily nalfurafine (50 ug/kg), nalfurafine is still able to block GNTI-induced itch (Fig. 5F). These findings suggest that the ROS-induced KOR inactivation is restricted to a subset of KOR responses. The basis for this tissue selectivity has not been established but presumably results from differences in JNK/PRDX6 expression or receptor regeneration rate.

## Discussion

The principal conclusions of this study are that activation of the JNK/PRDX6 pathway by nalfurafine and nalmefene results in norBNI-like KOR inactivation. The neuroimaging and behavioral pharmacological results provide substantial validation of the novel JNK signaling and ROS production mechanism proposed to be responsible for the long duration of KOR inactivation, which may be a general regulatory mechanism of Gi/o coupled receptor function. Because KOR inactivation is long-lasting, daily microdosing of either nalfurafine or nalmefene allows slow accumulation of inactivation of KOR necessary to block the pro-depressive effects of elevated dynorphin tone evident during stress exposure and abstinence in opioid-dependent mice. Estrogen regulation of the JNK/PRDX6 signaling pathway was confirmed in female mice, and progesterone treatment may be useful in enhancing treatment efficacy in premenopausal women. Most importantly, this study suggests that the KOR-inactivating medications nalfurafine and nalmefene, which have demonstrated safety in humans, may have potential advantages over conventional competitive antagonists in the treatment of depression and addiction disorders.

The results provide additional validation of the concept that KOR activates three different signaling pathways: Gβγ for membrane-delimited regulation of ion channel conductance; GRK3/β-arrestin regulation of p38 MAPK mechanisms controlling mood; and JNK/PRDX6 controlling Gαi coupling to KOR. KOR ligands differ in their efficacies at each pathway, and functional selectivity needs to be considered in at least these three dimensions. The long duration of norBNI was initially thought to be caused by slow pharmacokinetic clearance; however, an alternative explanation was suggested when norBNI was found to stimulate the phosphorylation of cJun N-terminal kinase (JNK) in KOR-transfected HEK293 cells and mouse brain (22). The long duration of norBNI action was prevented if JNK was pharmacologically inhibited in vivo or if the *JNK1* gene isoform was deleted (25). Additionally, the pharmacokinetic explanation was excluded by a receptor occlusion experiment (22), and the duration of action of a broad range of KOR antagonists was tightly correlated with their abilities to stimulate JNK phosphorylation in the spinal cord (25).

NorBNI enhances the association of KOR with Gβγ and PRDX6 which is known to stimulate NADPH generation of ROS through its phospholipase activity (46). ROS can oxidize sulfhydryl groups on exposed cysteines (47) and the amino-terminal cysteine residue in Gαi is reversibly palmitoylated. This palmitoylation controls the binding orientation of the Gαi in the receptor signaling complex (48). and norBNI generation of ROS depalmitoylates the Gαi (27). This depalmitoylation and the resulting inactivation are irreversible, requiring full resynthesis of the receptor (over 3 weeks) to recover the KOR response (22). Our results using the novel oROS sensor showed that ROS is generated by VTA KOR activation with both nalfurafine and nalmefene in ex vivo slice and in vivo fiber photometry and that KOR inactivation can be perturbed by blocking the JNK/PRDX6 pathway. The mechanism described here for KOR inactivation due to downstream ROS production also controls mu-opioid receptors (MOR). It may contribute to the tolerance seen when using some MOR agonists (25).

There is now a large body of preclinical literature suggesting that KOR antagonists may prove useful in the treatment of the anhedonic component of many mental disorders. NorBNI and JDTic, long-lasting KOR antagonists, have been shown to block stress-induced drug-seeking and drug self-administration (12, 49, 50). Dynorphin is elevated in animal models of stress behaviors and in the CSF of persons experiencing psychosis (51). In this study, we show the ability of nalfurafine and nalmefene to block stress-cue-paired aversion. In the odor-swim stress pairing, there is a full block of odorant aversion, consistent with what has been shown with a single large dose of norBNI (42). In the odorant-opioid withdrawal pairing, KOR inactivation caused a partial block during acute withdrawal and a complete block during protracted withdrawal. Acute opioid withdrawal includes a profound contribution of a hyperactive sympathetic nervous system, and the physical symptoms can be ameliorated by α2A adrenergic receptor agonists clonidine or lofexidine (52). Naloxone precipitation is not identical to spontaneous withdrawal from opioids - it is more intense and acute; however, the symptoms are qualitatively similar (53). Although we are ultimately interested in determining the role of dynorphin in humans experiencing opioid abstinence, the conditioning paradigm in mice works better if the cue and the stimulus are discrete and temporally associated. The results suggest that during the protracted withdrawal phase, the dysphoric effects are largely mediated by residual hyperactivity of the dynorphin/KOR system. Therefore, KOR antagonists may have therapeutic utility to promote stress resilience during protracted abstinence in humans.

Theoretically, KOR-inactivation by nalfurafine or nalmefene confers several potential advantages as medications. First, these two medications have histories of safety and tolerability in human use. Nalfurafine has been approved in Japan as an antipruritic in uremic itch (35). Nalmefene is used as an alternative to naloxone and naltrexone in the treatment of alcohol use disorder and to reverse respiratory depression during acute opioid overdose (54). Slow accumulation of inactivation would allow the titration of effect, and slow recovery would allow stable antagonism. In contrast, competitive antagonists would need to be taken at supersaturating doses to prevent dynorphin activation of KOR. Dynorphins have very high affinities and receptor efficacies; they may need only occupy a fraction of the receptors to produce a maximal effect (55). The results presented in this study highlight the potential of leveraging functional receptor selectivity insights for therapeutic advantage. With the identification and characterization of these three signaling pathways at KOR, we can expand beyond the traditional idea of partial agonism to generate novel medications to promote stress resilience.

## Author Contributions

M.V.R., K.K., and C.C. contributed equally to data collection and analysis. Conceptualization: C.N., B.L., C.C. Methodology: C.N., B.L., A.B., C.C., Investigation: C.N., M.V.R., K.K., C.C., J.D.L Visualization: C.N., M.V.R., K.K., J.D.L. Funding acquisition: A.B., C.C. Project administration: C.C. Writing – original draft: C.N., C.C., Writing – review and editing: C.N., M.V.R., K.K., J.D.L, B.L., A.B., C.C.

## Acknowledgments

We thank Lucia Aballay, Varun Mehta, Anabelle Shen, and Claire Lee for assistance with microinjection and data acquisition.

## Diversity, equity, ethics, and inclusion

One or more of the authors of this paper identify as an underrepresented ethnic minority in science. One or more of the authors of this paper identify as a member of the LGBTQ+ community.

## Data and materials availability

All data are available in the main text or the supplementary materials.

