## Supplemental Figures for "G protein Inactivation as a Mechanism for Addiction Treatment"

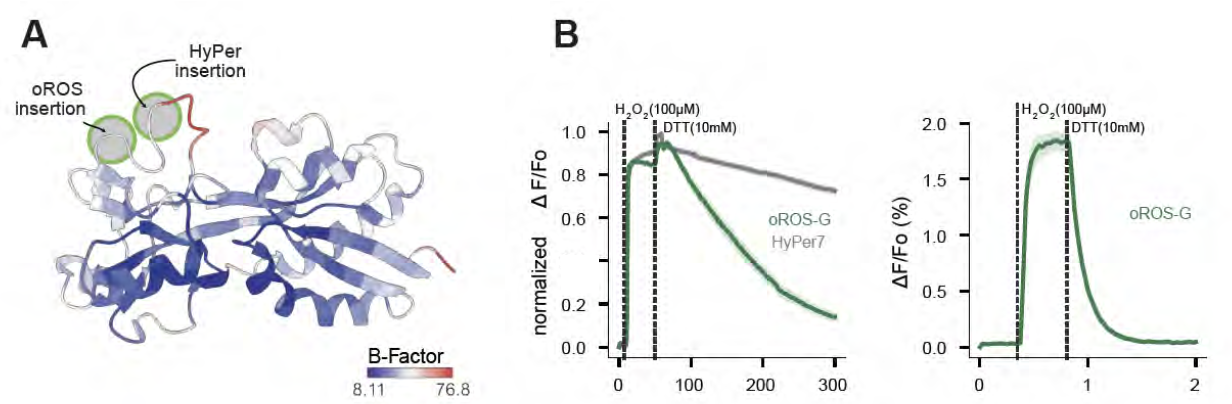

**Fig. S1: Enhanced sensitivity and kinetics of the oROS sensor. (A)** Crystal structure of reduced and oxidized forms of Regulatory Domain (RD) of Escherichia coli OxyR. C-C pair labeled in yellow. Red indicates the fluorescent protein insertion loop for HyPer sensors, and Blue indicates the newly identified fluorescent protein insertion site for oROS sensors. (**B)** Representative reduction kinetics of oROS-G and HyPer7 after 100 µM H_2_O_2_ stimulation followed by media wash (n>100 cells per sensor). HEK293 expressing oROS-G were stimulated with 100 µM H_2_O_2_ followed by media wash.

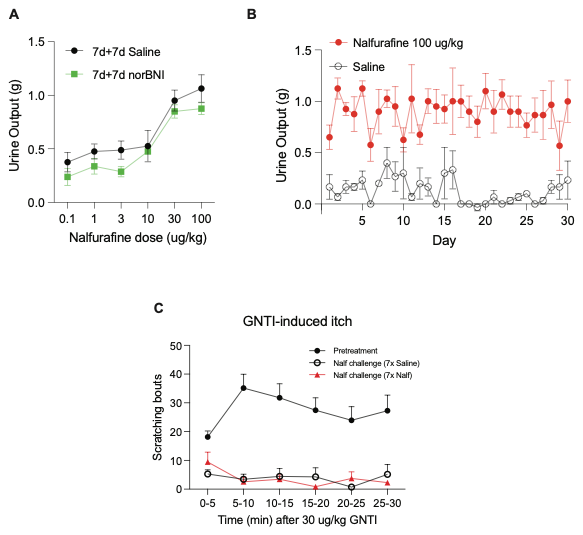

**Fig. S2: KOR inactivating agents demonstrate short-acting antagonism in specific tissues. (A)** Nalfurafine was given to mice each day for 6 days at varying doses with pretreatment of either saline or low-dose norBNI (1 mg/kg) for 7 days and 7 days of recovery. No differences were observed between saline and norBNI repeated dose pretreatment. Doses of nalfurafine were given daily, and tolerance to its diuretic action was not observed.  (**B)** Mice treated daily with 100 mg/kg nalfurafine (n=4) showed no tolerance to the diuresis effect of nalfurafine. Saline (n=3) had no effect on diuresis. (**C)** GNTI-induced itch data from Fig. 4F displayed in 5 min bins.
